# A 29-year retrospective analysis of koala rescues in New South Wales, Australia

**DOI:** 10.1101/2020.04.19.048967

**Authors:** Renae Charalambous, Edward J. Narayan

## Abstract

The koala (*Phascolactos cinereus*) is currently listed by both the IUCN and the Australian Governments’ Threatened Species Scientific Committee as vulnerable to extinction with a decreasing population trend. This listing can be attributed to climate change and its impact on ecosystems, and anthropomorphic environmental change due to extensive land clearing and habitat fragmentation. These have both been proven to induce stress, which influences the onset of disease. This novel study performed a retrospective analysis whereby records for 12,543 wild, rescued koalas in New South Wales (NSW), Australia were studied in order to determine trends in koala sightings, clinical admissions and injury diagnoses over a period of 29 years (1989-2018). Results indicated that between all three study locations (Port Stephens, Port Macquarie and Lismore), the most common reason koalas were admitted into care was because of disease, the most common disease diagnosed was signs of chlamydia, and the most common outcome for koalas admitted into care was released. At Port Stephens, mature and female koalas were diagnosed with a disease more than any other age or sex, while juvenile and male koalas were released (back into the wild) more than any other age or sex. Additionally, there were fewer koalas with a disease and fewer koalas released in Port Stephens as each year progressed. At Port Macquarie, mature and male koalas were diagnosed with a disease more than any other age or sex, while juvenile and female koalas were released more than any other age or sex. Additionally, there were more koalas with a disease and fewer koalas released in Port Macquarie as each year progressed. At Lismore, adult and female koalas were diagnosed with a disease more than any other age or sex, while joey and male koalas were released more than any other age or sex. Additionally, there were more koalas with a disease and fewer koalas released in Lismore as each year progressed. Determining trends in clinical admissions and diagnosis over such a substantial period of time is an important factor in preventing the continuing decline of koalas throughout Australia, and in particular NSW. It is important to note that there are cultural differences between koala rescue groups in the three study locations (Port Stephens, Port Macquarie and Lismore). These differences may be reflected in the outcomes of koala patients as each group are driven by their own management team. It is essential that any further decline of koala populations is prevented, however this can only be achieved through informed recommendations through research studies such as these. These recommendations should lead to government legislation which can provide stronger protection to koala habitat.

## Introduction

In theory, Australia should have relatively few conservation concerns; it’s national population density is low (~3km^−2^) by global standards (~50km^−2^), most of the continent remains sparsely settled and little modified, and the nation is relatively affluent[1]. However, since 1788, 30 mammal species endemic to Australia have become extinct, with 55 others experiencing a worsened conservation status[2]. This statistic is primarily attributed to the fact that humans alter areas of pristine habitat that is rich with biodiversity, so as to accommodate their rapid population growth within coastal areas of Australia[2].

One of Australia’s most iconic animal species is the koala, and this species has not been exempted from the aforementioned conservation concerns. Koalas living in the North-East of Australia have experienced a 40% clearance of ideal habitat and a population decline of 80% since 1990[3]. It is unknown how many koalas remain in the wild throughout all of Australia as population parameters vary. The Australian Koala Foundation estimate that there could be less than 80,000 wild koalas remaining in Australia[4], whereas a report by the Chief Scientist of Australia estimate this figure to be around 330,000[5]. Despite being listed as “vulnerable to extinction” by both the IUCN in 2014, and the Australian Governments’ Threatened Species Scientific Committee in 2012, the conservation status for the koala remains highly contentious primarily due to the uncertainty of population parameters[6],[7]. Furthermore, with respect to the eligibility of koalas against the IUCN criteria, both drought and bushfires were listed as the predominant threats associated with this species[6]. Contrary to this, there is now mounting evidence to suggest that extreme environmental factors alone are not the biggest threat affecting populations of koalas in Australia[8]. Human led changes to Australia’s natural landscape both inland and along the coast have created ecological problems for koalas[8]. For instance, human population growth is highest throughout the East Coast of Australia, which amplifies land clearance and competition for suitable habitat between koalas and humans[8]. Due to the challenge of accurately estimating their population size, and the increased human induced risks associated with their remaining population, it is absolutely necessary that the conservation status for koalas be re-evaluated, and efforts are made to accurately measure the number of koalas remaining throughout Australia.

As outlined in a study of koalas living in Queensland (QLD), Australia, the biggest threats affecting populations of this species derive from human population growth and global climate change [8]. Due to the removal of suitable habitat by tree clearing to cater for a growing human population, there are a high proportion of koalas who present to clinical care with injuries consistent with motor vehicle trauma and trauma stemming from animal attacks[9]. Between 1997 and 2013 in SE QLD, 15.5% of koalas admitted into clinical care were diagnosed with motor vehicle trauma, and 5.2% of koalas admitted into clinical care were diagnosed with trauma stemming from animal attacks[9]. Of particular concern is that these koalas had no symptoms of underlying disease, suggesting potential for impact on local population growth by healthy, breeding koalas being prematurely removed[9]. Due to increased pressure from global climate change and the extension of human settlement across suitable koala habitat, there is a high proportion of koalas who present to clinical care with symptoms of chlamydia disease[9]. Between 1997 and 2013 in SE QLD, 66.3% of koalas admitted into clinical care were diagnosed with chlamydia disease or presented signs consistent with chlamydia disease[9]. Infectious pathogens that rely on frequency-dependent transmission, such as chlamydia disease can influence population dynamics by increasing mortality from wasting and blindness, and decreasing population recruitment through impairment of reproduction[9]. However, studies have found that diseases such as chlamydia will only play a detrimental role within a population when other extrinsic factors inducing physiological stress are present, or extensive periods of droughts and clearance of native vegetation for human use [8, 9].

This study is a retrospective analysis whereby records for 12,543 wild, rescued koalas in New South Wales (NSW), Australia were studied in order to determine trends in koala sightings, clinical admissions and injury diagnosis over a period of 29 years (1989-2018). This study is important as a significant lack of published data specific to the relationship between stress and koalas exists. Furthermore, this study aims to contribute valuable information relating to the current conservation status of koalas as there is a vast amount of uncertainty relating to population parameters. Performing a retrospective analysis of koala sightings, clinical admissions and injury diagnoses will provide an educated recommendation to update the conservation status to one which is more representative.

It is hypothesised that there will be a difference between the prognosis and outcome of koalas based on:

1. year admitted into care (1989-2018)
2. location admitted into care (Port Stephens, Port Macquarie and Lismore; see methods for GPS information)
3. sex (male and female)
4. age (adult, joey, juvenile and mature)

## Methods

### Approval

All research was done in accordance with the following relevant guidelines and regulations. All current research with NSW koalas was formally approved by the Western Sydney University Animal Care and Ethics (ACEC) Committee approval number (Protocol number: A12373).

### Data Analysis

Admission forms and hospital records for 12,543 wild koalas reported or admitted to one of three locations (Port Stephens Koalas in Port Stephens, Port Macquarie Koala Hospital in Port Macquarie and Friends of the Koala in Lismore) over a 29-year period (1989-2018) were collected and analysed.

Port Stephens Koalas is located at 562 Gan Gan Road, One Mile (GPS Coordinates: - 32.763792, 152.115904). At maximum capacity, the hospital can hold 20 koalas at one time, however when koala numbers exceed capacity, long-term carers are able to care for these animals in a foster home setting until they are back to good health. Port Macquarie Koala Hospital is located within the Roto House Historical Site on Lord Street, Port Macquarie (GPS Coordinates: −31.442102, 152.919167). At maximum capacity, the hospital can hold 100 koalas at one time. Friends of the Koala is located at 23 Rifle Range Road, East Lismore (GPS Coordinates: −28.820714, 153.302499). At maximum capacity, the hospital can hold 25 koalas at one time, however when koala numbers exceed capacity, long-term carers are able to care for these animals in a foster home setting until they are back to good health.

The information recorded included each patient’s sex, age, suburb found, prognosis, prognosis sub-category, outcome, and year admitted. The information for all 12,543 koalas were recorded in Microsoft^®^ Excel, then were analysed in IBM SPSS Statistics^®^. Two tests were run for each variable in the hypothesis, with the aim of summarising the data to determine patterns. These tests included a descriptive statistical analysis and a generalised linear model. Age, sex, place and year (independent variables) were all plotted against both the admission prognosis and outcome (dependant variables), and the results were displayed as column graphs. Raw data for descriptive statistics was transformed into proportions to enable comparisons, and this made it possible to successfully plot age, sex, place and year against prognosis and outcome.

For each koala patient, the suburb where they were found was recorded to determine distribution trends. ArcGIS was used to map the distribution of koalas admitted to all three koala hospitals (Port Stephens Koalas, Port Macquarie Koala Hospital and Friends of the Koala). Postcodes and the number of koalas admitted within each postcode were transcribed into comma-separated documents (CSV) on Microsoft^®^ Excel. The CSV sheets were then uploaded separately as a base layer on ArcGIS. Once uploaded, a dot distribution map was generated and koala “hot zones” were attained.

Finally, using Microsoft^®^ Excel, sighting and admission records for all 12,543 koalas were used to graph the frequency of both prognosis and disease. Percentages for all koalas were calculated in order to determine prognosis, disease and outcome, and the percentages were then used to create three column graphs, with the data displayed in descending order.

It is important to note that due to the stochastic nature of admissions into clinical care based on altered population density, the number of koalas admitted into each hospital, different management strategies by each koala hospital and the available temporal data was not uniform at each location. As a result of this, proportional data was used when comparing locations while the majority of analyses focused on trends within individual sites.

## Results

### Frequency and locations of koala rescues

The suburbs where koalas were most frequently found prior to being reported as sighted or admitted into clinical care at Port Stephens Koalas were Salamander Bay (335 koalas), Anna Bay (276 koalas) and One Mile (195 koalas). At the Port Macquarie Koala Hospital, the most frequent suburbs were Port Macquarie (430 koalas), Armidale (14 koalas) and Limeburners Creek (13 koalas). For Friends of the Koala, the most frequent suburbs were Goonellabah (1,219 koalas), Lismore (1,004 koalas) and Wyrallah (444 koalas) [Figure 1.0 A, 1.0 B & 1.0 C; Also see supplementary file 1 for proportion and frequency of koala rescues at each site].

**Figure 1.0:**
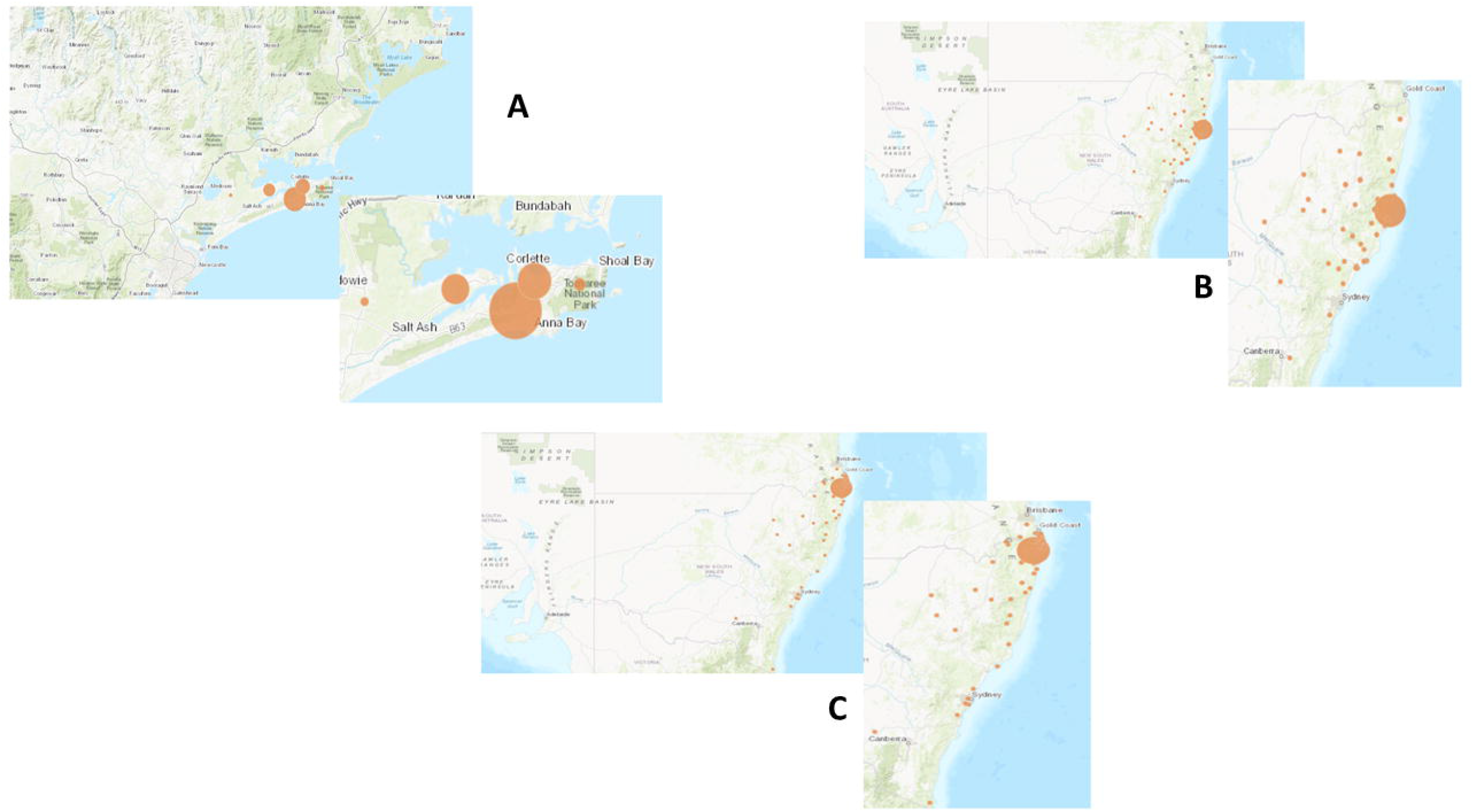
A: Distribution trends displayed as hot zones for koalas admitted into care at (A) Port Stephens Koalas, (B) Port Macquarie and (C) Friends of the Koala between 1989 and 2018.

### Prognosis and outcome of koala rescues

Each koala was given a prognosis after being reported as sighted or admitted into clinical care. These included appearing healthy on assessment, being attacked by cattle, being collared for tracking, being diagnosed with a disease, being dead on arrival, being attacked by a dog, being caught in a fire, being harassed by humans, being hit by a car, being an orphan, being attacked by a snake, having an unknown prognosis, or being displaced in an unsuitable environment. Of all 12,543 koalas over the three locations, 34.5% were recorded as diseased, 24.4% were recorded as having an *unknown prognosis*, and 16.2% were recorded as *appeared healthy* (Figure 2.0).

**Figure 2.0:**
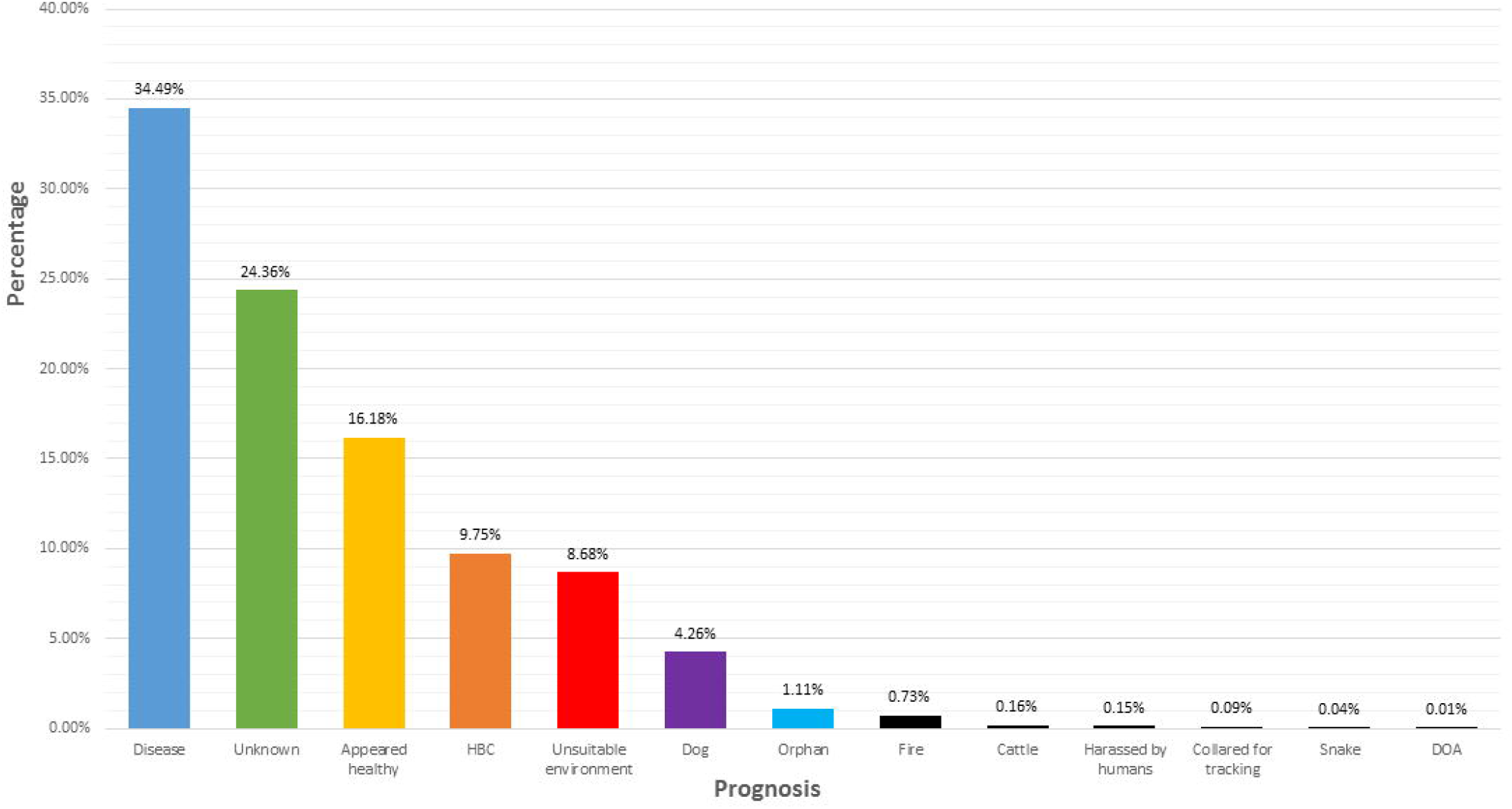
The prognosis that was given to each wild rescued koala patient (n=12,543), displayed as a percentage, when they were admitted to one of three locations (Port Stephens Koalas, Port Macquarie Koala Hospital and Friends of the Koala) between 1989 and 2018.

Of the diseases recorded, included was having signs of chlamydia, infection, poor body condition, damage to one or many organs, injuries to one or both eyes, old age, experiencing trauma to the head, dehydration, tick infestation, koala retrovirus, injuries to one or many claws, and injury to one or both legs. Of all the koalas who were recorded as having a disease, the most common was *signs of chlamydia* (51.5%), followed by *infection* (24.2%), and *poor body condition* (10.1%) (Figure 3.0).

**Figure 3.0:**
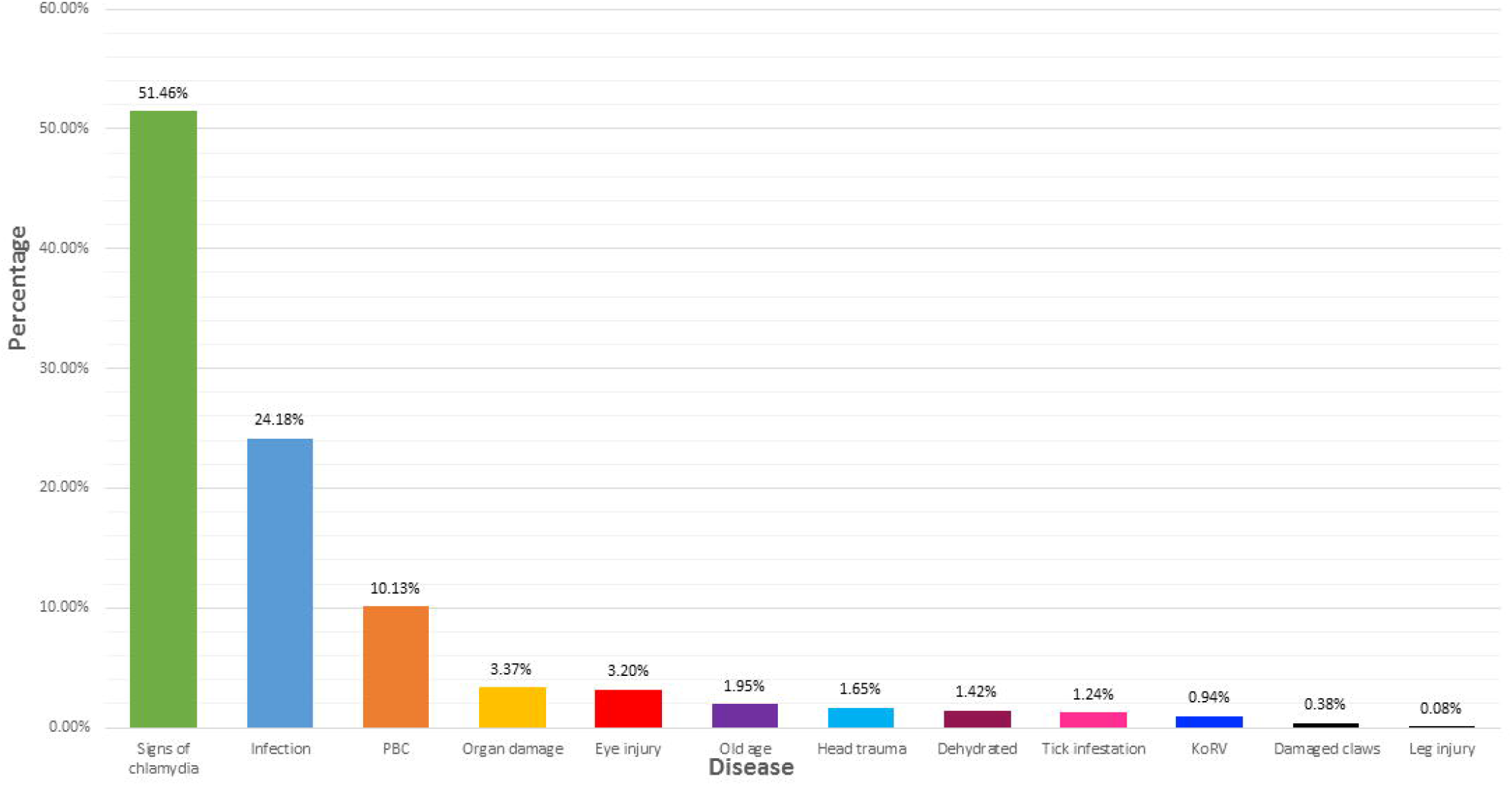
The disease diagnosis that was given to each wild rescued koala patient (n=3,941), displayed as a percentage, when they were admitted to one of three locations (Port Stephens Koalas, Port Macquarie Koala Hospital and Friends of the Koala) between 1989 and 2018.

Each koala was given an outcome after being reported as sighted or admitted into clinical care. These included being released into the wild, being euthanised, being dead on arrival, dying while in care, being trapped and relocated to a new area, unknown, transferred to a different captive facility, still in care, and koala escaped from care/self-release (note, some outcomes included providing advice to the member of public, recording a sighting of a koala, unable to capture, and member of the public would not hand the koala over to the carer; although these koalas were not admitted into care, it is still relevant for understanding population dynamics). Of the 12,543 koalas over the three locations, 20.7% were released, 17.4% were *euthanised*, and 17.3% required *advice only* (Figure 4.0).

**Figure 4.0:**
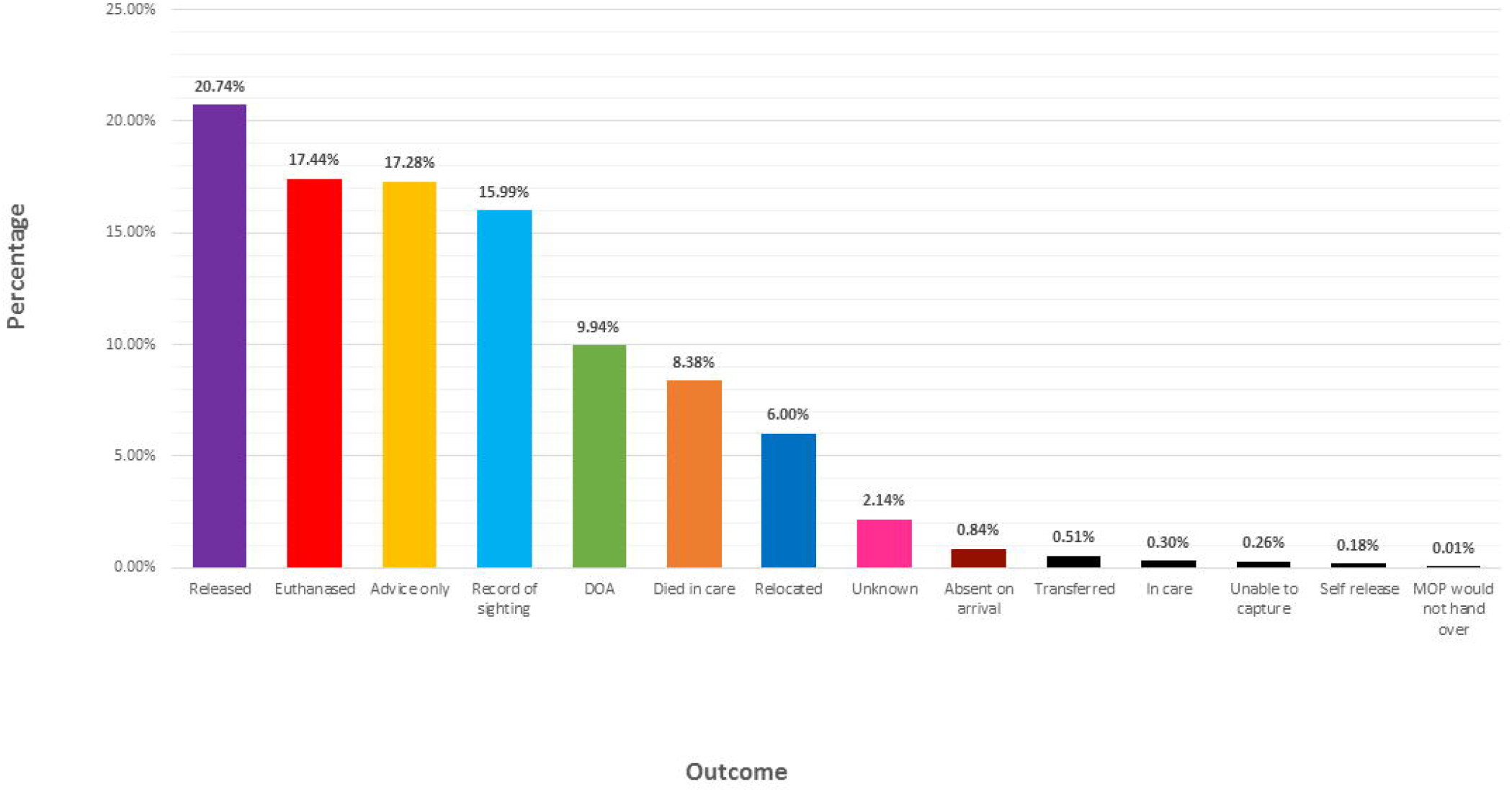
The outcome that was given to each wild rescued koala patient (n=12,543), displayed as a percentage, after they were admitted to one of three locations (Port Stephens Koalas, Port Macquarie Koala Hospital and Friends of the Koala) between 1989 and 2018.

### Comparing prognosis and outcome with age, sex, location and year

*Prognosis categories* include appearing healthy on assessment, being attacked by cattle, being collared for tracking, being diagnosed with a disease, being dead on arrival, being attacked by a dog, being caught in a fire, being harassed by humans, being hit by a car, being an orphan, being attacked by a snake, having an unknown prognosis, or being displaced in an unsuitable environment (some prognosis categories were excluded as they were statistically non-significant).

*Outcome categories* include being absent on arrival, providing advice to the member of public, died in care, dead on arrival, euthanised, still in care, member of the public refusing to admit the koala to the hospital, record of sighting, released into the wild, relocated to a new location, released themselves/escaped from clinical care, transferred to another caring facility or organisation, unable to capture, and unknown (some outcome categories were excluded in figures as they were statistically non-significant. Furthermore, some outcomes involved koalas not physically being admitted into care, however, were still included as it is still relevant for understanding population dynamics).

### Age compared with prognosis and outcome

Age categories for koalas were grouped according to life stages, and they included joey, juvenile, adult, mature and unknown (see discussion for an in-depth review of life stages). The most common prognosis between all three locations was *disease* (34.4%), and the age group that was affected the most by this prognosis was *mature* koalas (42.2%). The most common outcome between all three locations was *released* (20.7%), and the age group that was affected the most by this prognosis was *mature* koalas (45.8%).

### Sex compared with prognosis and outcome

Sex categories for koalas include male, female and unknown (the unknown sex is due to koala sightings, as sex cannot be confirmed from a distance). The most common prognosis between all three locations was *disease* (34.4%), and the sex category that was affected the most by this prognosis was considerably even with *female* koalas (38.9%), and *male* koalas (38.3%). The most common outcome between all three locations was *released* (20.7%), and the sex category that was affected the most by this prognosis was *female* koalas (33.0%).

### Location compared between prognosis and outcome

The prognosis that occurred the most between all three locations (Port Stephens, Port Macquarie and Lismore) was *disease* (34.4%), and the location category that was affected the most by this prognosis was *Lismore* (37.1%). The outcome that occurred the most between all three locations was *released* (20.7%) and the location category that was affected the most by this outcome was *Port Macquarie* (71.4%).

### Year compared between prognosis and outcome

Year categories ranged from 1989-2018. The prognosis that occurred the most between 1989-2018 was *disease* (34.5%), and the year that was affected the most by this prognosis was *1992* (56.0%). The outcome that occurred the most between 1989-2018 was *released* (20.7%) and the year that was affected the most by this outcome was *1989* (50.7%).

## Discussion

The aim of this study was to perform a retrospective analysis whereby records for 12,543 wild, rescued koalas in NSW were studied in order to determine trends in koala sightings, clinical admissions and injury diagnosis over a period of 29 years (1989-2018). Three major koala hospitals within NSW were sampled (Port Stephens Koalas, located in Port Stephens, Port Macquarie Koala Hospital, located in Port Macquarie, and Friends of the Koala, located in Lismore) whereby a total of 12,543 records for koalas were collected.

Between the 3 major koala hospitals, 12,543 koalas were either reported as a sighting or admitted into care (Figure 1.0 A, 1.0 B & 1.0 C). Identifying habitat preference is central to conservation, and it is important when trying to understand what the threats that have led to their admission into clinical care are. Habitat is a species-specific concept defined as “the resources and conditions present in an area that produce occupancy”[10]. For koalas, a home-range should include habitat resources that provide the necessary food, shelter and space requirements for reproduction, survival and movement between trees[10]. Survival and population growth is heavily dependent on suitable home-ranges, and lack thereof has direct implications for the risk of extinction[10]. The major factor defining a suitable home-range for a koala population depends on their preference for a relatively small number of eucalypt species that are able to provide a nutrient-rich foliage[11]. Evidence suggests that foliar chemistry can explain differences in nutritional value and subsequent food selection by koalas between and among eucalyptus species[12]. Studies have shown that there is a higher-frequency of visitation by koalas compared to other species to eucalypts such as the Tasmanian blue gum (*Eucalyptus globulus*) and manna gum (*Eucalyptus viminalis*)[13]. This is likely due to these eucalypt species containing higher concentrations of nitrogen and lower concentrations of plant secondary metabolites, and this suggests that foliar chemistry may influence koala distribution and abundance[13]. Despite all 3 study locations containing climatically suitable home ranges for koalas, their populations are experiencing a steep decrease, and many are admitted into clinical care[6]. This can be attributed to increased spatial competition between koalas and humans with anthropogenic activity resulting in reduction and fragmentation of suitable koala habitat[14, 15]. In particular, according to the NSW Government, regional areas such as Lismore are more susceptible to habitat fragmentation, as they have experienced an influx of tourists due to excellent tourism infrastructure, facilities, and proximity to the natural environment. Compared to areas such as Port Stephens and Port Macquarie who are not classified as regional areas, Lismore is a prime location for animal agriculture and crop growth.

It is a clear commonality that literature suggests koala populations are declining as a direct result of human activity, most notably habitat destruction and fragmentation[3]. Any disturbance to an animal’s home range activates the physiological stress response[15], and if the stressor/s is not mitigated, the excessive production of glucocorticoids can leave the animal with a compromised immune system[8]. It is therefore not surprising that of the 12,543 koala’s ether recorded as being sighted or admitted into care through 1989-2018, 34.5% presented with a disease (Figure 2.0). Of the 34.5% of koalas that presented with a disease, 51.5% of them were diagnosed with chlamydia or displayed signs of chlamydial infection (Figure 3.0). *Chlamydia pecorum* is an infectious bacterial pathogen that operates as a significant threat to koala conservation[16]. Chlamydia is primarily a sexually transmitted infection in koalas, however there is anecdotal evidence for vertical transmission (transmission of a pathogen from mother to baby either during birth, or after birth)[16]. Efforts to understand this disease have found ocular infections of chlamydia can lead to debilitating blindness, while urogenital tract infections of chlamydia can lead to cystitis and/or ascending infections of the reproductive tract and sterility[17]. In the wild, chlamydia in koalas can be identified through red, inflamed eyes and a brown, wet bottom[18]. Koalas infected with this disease often starve to death as the ocular infection creates proliferative inflamed conjunctival tissue that can grow over the cornea of the eye, rendering the koala blind and unable to find appropriate food or shelter[19]. Depending on the stage of chlamydia, koalas admitted into clinical care can be administered with antibiotics as a treatment for the disease, although keeping in mind that this can adversely affect the gut microflora and health of the animal[18]. A late diagnosis of chlamydia may require euthanasia on the grounds of welfare, as the disease is incredibly painful for the infected koala[18].

This retrospective analysis identifies some important trends in koala sightings, clinical admissions and injury diagnosis (Figures 1.0 – 4.0). In most instances, adult and mature aged koalas were more susceptible than joey and juvenile aged koalas to disease, and female koalas were more susceptible than male koalas to disease. The variation in sex and age specific selection pressures can have a fundamental influence on animal survival[20]. This is due to the fact that fitness requirements differ between sexes and among age classes in distinctive ways[20].

When koalas are born, they are roughly 2cm long and remain blind and hairless in their mothers’ pouch feeding off milk from a teat; at this stage, they are referred to as either a joey or a pinkie [4]. At about 6 months to 1 year old, the koala exists the mothers’ pouch and weans off of milk and on to leaves[4]. To assist this transition, the mother will produce a substance called “pap”, which is a micro-organism necessary for making digestion for her juvenile koala possible[4]. Once they are a year old, they will prepare to leave their mother, and at this stage they are referred to as a juvenile[4]. At about 15 months of age, female koalas are sexually mature, and at about 2 years of age, male koalas are sexually matured. At this stage, they are referred to as an adult[4]. From 5-7 years and above, koalas’ transition from an adult to a mature koala[4]. At both the adult and mature stage of life, survival becomes harder for koalas as they no longer have their mother to protect them from predators, and to provide food and shelter. As an adult, koalas are frequently moving with the potential to cross over roads with cars and enter landscapes with dogs and other predators. These reasons are the likely contributors to our results indicating that adult and mature aged koalas were more likely than joey and juvenile aged koalas to experience disease among other identified stressors.

Female koalas begin breeding when they are at 15 months of age, and generally produce one offspring per year[4]. During and after gestation are both a nutritionally demanding time for a female koala, as they are required to not only feed themselves, but also their young[4]. This can become extremely stressful if resources become scarce, and this could easily take a toll on the female koala during not only a 35-day gestation period, but the remaining 330 days that they are responsible for their young. Unlike female koalas, at no stage in their lives are male koalas responsible for another koala, young or old. The higher mothering expectations on females over males are likely why our results indicated that female koalas were more likely than male koalas to experience disease among other identified stressors.

The number of koalas being admitted into clinical care increased progressively over each year. Despite this, the proportion of koalas released decreased across all three study locations. During 1990s, the Australian population was estimated at 17,041,431 people, and by 2018, was estimated at 24,772,247 people. The difference of 7,730,816 people over a 28-year period is dramatic especially as it is argued that the environment has reached the upper limits of human population growth[21]. At the very least, the East-Coast of Australia, where koalas occupy their home ranges, are being encroached on as the number of humans increase. The presence of humans is closely tied to koala stress, and this is likely why our results indicated that more koalas were being admitted into clinical care over each year. These trends arise despite early detection as a result of growing awareness of koalas in Australia. Similarly, as facilities such as Port Stephens Koalas, Port Macquarie Koala Hospital and Friends of the Koala receive slight increases in funding through donations and government grants, so too do their facilities and access to treatment improve.

It is integral to formulate a response to the increasing threats to koalas. One management response to decreasing suitable habitat for koalas is translocation. Translocation is a wildlife management tool employed to establish, re-establish or augment wildlife populations, particularly in response to intentional habitat destruction, to minimise human-wildlife conflicts, and to reduce densities of over-abundant species[22]. Previously, translocation has been used in the management of koala populations to reduce the effect of high-density populations on forest habitats and lessen the risk of density-related animal welfare issues[22]. One study aimed to research the success of translocation on koala populations, as it is a possible conservation option for koala populations currently experiencing a steep home range decline. Between 1997 and 2007, over 3,000 koalas were captured, sterilised and translocated to other home-ranges in mainland Australia[22]. Results of the translocation revealed densities of <0.4 koalas/hectare, despite release densities of 1.0 koalas/hectare[22]. Radiotracking indicated that the low densities could be attributed to a 37.5% mortality of translocated individuals within the first 12 months, post release[22]. Evidence suggests that koalas do not exhibit high survival rates in response to translocation, and therefore, these animals remain in their compromised home ranges [16]. With such high mortality rates associated with translocation, it could be argued that a more appropriate response would be the augmentation of human behaviour, selecting alternate areas for investment and development.

## Conclusion

Through this novel retrospective analysis of 29-years’ worth of data of koalas in NSW, Australia, it is demonstrated that koalas are in a dire state requiring stronger conservation and management. The aim of this study was to perform a retrospective analysis whereby records for 12,543 wild, rescued koalas in NSW were studied in order to determine trends in koala sightings, clinical admissions and injury diagnosis over a period of 29 years (1989-2018). The first hypothesis, that there will be a difference between the prognosis and outcome of koalas based on year admitted into care (1989-2018), can be accepted. Until 2005, the most frequent occurring prognosis was disease, and the most frequent occurring outcome was released. From 2005 until 2012, the most frequent occurring prognosis fluctuated between disease and appearing healthy (admission was mostly due to trauma from factors such as dog attacks or motor vehicle collision, as well as possible cases of trauma from bushfires), and the most frequent occurring outcome was advice only. After 2012, the most frequent occurring prognosis continued to fluctuate between disease and appearing healthy, and the most frequent occurring outcome was record of sighting. The prognosis and outcome of koalas admitted into care continually changed between the years 1989 and 2018, and this clearly reflects societal awareness of koala presence as well as the increased prevalence of disease in koala populations over a 29-year period. The second hypothesis, that there will be a difference between the prognosis and outcome of koalas based on location admitted into care (Port Stephens, Port Macquarie & Lismore), can be accepted. The most frequent occurring prognosis for koalas admitted into care within Port Stephens and Port Macquarie was based on unsuitable habitat, whereas in Lismore disease was most common. The most frequent occurring outcome for koalas admitted into care within Port Stephens and Port Macquarie was released, whereas for koalas admitted into care within Lismore was advice only. The prognosis and outcome of koalas admitted into care continually changed between the three locations, and this clearly reflects how environment change affects koala populations differently based on individual locations over a 29-year period. The third hypothesis, that there will be a difference between the prognosis and outcome of koalas based on their sex (male & female), can be rejected. For both male and female koalas admitted into care, the most frequent occurring prognosis was disease, while the most frequent occurring outcome was released. This clearly demonstrates that both male and female koalas are equally susceptible to disease and equally likely to be released after being admitted into clinical care. The fourth hypothesis, that there will be a difference between the prognosis and outcome of koalas based on their age (adult, joey, juvenile & mature), can be accepted. For adult koalas, the most frequent occurring prognosis was disease, while the most frequent occurring outcome was advice only. For joey koalas, the most frequent occurring prognosis was disease, while the most frequent occurring outcome was released. For juvenile koalas, the most frequent occurring prognosis was disease, while the most frequent occurring outcome was released. Finally, for mature koalas, the most frequent occurring prognosis and outcome were disease released respectively. The prognosis and outcome of koalas admitted into care remained the same for joey, juvenile and mature aged koalas, but differed for adult aged koalas, indicating that koala populations are affected differently based on their age bracket. In conclusion, this retrospective analysis determined trends in clinical admissions and diagnosis for koalas admitted into care within NSW, Australia over a 29-year period. Further detailed investigation is warranted on the synergistic nature of anthropogenic induced environmental trauma especially land clearance and prolonged climatic variability such as bushfires and droughts towards aggravating infectious diseases and increasing the risks of koalas becoming exposed to acute environmental trauma such as heat stress, and victims of dog-attacks and vehicle trauma. This enables any steps to mitigate the further decline of koala populations to be better informed, with the implementation of scientific based management strategies for effective koala conservation.

## Acknowledgements

RC conducted Master of Research under the principal supervision of EN. Thanks to the veterinary staff, nurses and volunteers of the Friends of the Koala, Port Macquarie Koala Hospital and Port Stephen Koala Hospital. Special thanks to the International Funds for Animal Welfare (IFAW) for supporting this research.

## Availability of data and materials

The Friends of the Koala, Port Macquarie Koala Hospital and Port Stephen Koala Hospital have all been duly acknowledged for providing clinical data that have been adequately presented in graphs and also available in the supplementary file.

## Authors’ contributions

RC conducted the literature collection and data analysis under the supervision of EN.

